# Integrating citizen science and field sampling into next-generation early warning systems for vector surveillance: Twenty years of municipal detections of *Aedes* invasive mosquito species in Spain

**DOI:** 10.1101/2025.08.04.668402

**Authors:** Roger Eritja, Isis Sanpera-Calbet, Sarah Delacour-Estrella, Ignacio Ruiz-Arrondo, Maria Àngels Puig, Mikel Bengoa-Paulís, Pedro María Alarcón-Elbal, Carlos Barceló, Simone Mariani, Yasmina Martínez-Barciela, Daniel Bravo-Barriga, Alejandro Polina, José Manuel Pereira-Martínez, Mikel Alexander González, Santi Escartin, Rosario Melero-Alcíbar, Laura Blanco-Sierra, Sergio Magallanes, Francisco Collantes, Martina Ferraguti, María Isabel González-Pérez, Rafael Gutiérrez-López, María Isabel Silva-Torres, Olatz San Sebastián-Mendoza, María Cruz Calvo-Reyes, Marian Mendoza-García, David Macías-Magro, Pilar Cisneros, Aitor Cevidanes, Eva Frontera, Inés Mato, Fernando Fúster-Lorán, Miguel Domench-Guembe, María Elena Rodríguez-Regadera, Ricard Casanovas-Urgell, Tomás Montalvo, Miguel Ángel Miranda, Jordi Figuerola, Javier Lucientes-Curdi, Joan Garriga, John Rossman Bertholf Palmer, Frederic Bartumeus

## Abstract

The spread of invasive mosquitoes *Aedes albopictus, Aedes aegypti* and *Aedes japonicus* in Spain represents an increasing public health risk due to their capacity to transmit arboviruses such as dengue, Zika and chikungunya among others. Traditional field entomological surveillance remains essential for tracking their spread, but it faces limitations in terms of cost, scalability and labor intensity. Since 2014, the Mosquito Alert citizen science project has enabled public participation in the surveillance through the submission of geolocated images via a mobile app, which are identified using AI in combination to expert validation. While field surveillance provides high accuracy, citizen science offers low-cost, large-scale real-time data collection aligned to open data managment principles. It is particularly useful for detecting long-distance dispersal events and has contributed up to one-third of the municipal detections of invasive mosquito species since 2014. This study assesses the value of integrating both surveillance systems to capitalize on their complementary strengths while compensating weaknesses in the areas of taxonomic accuracy, scalability, spatial detection patterns, data curation and validation systems, geographic precision, interoperability and real-time output. We present the listing of municipal detections of these species from 2004 to 2024, integrating data from both sources. Spain’s integrated approach demonstrates a pioneering model for cost-effective, scalable vector surveillance tailored to the dynamics of invasive species and emerging epidemiological threats.

**Simple Summary:** Spain, with its diverse ecological and climatic landscapes, represents a key area for the introduction and establishment of invasive mosquito species, posing threats to public health due to their capacity to transmit human diseases. Field entomological surveillance conducted by experts remains the primary tool for tracking their spread, though it is limited by geographical scales and cost factors. Advances in smartphone technology allows citizen science strategies of data collection such as submitting mosquito photos via the Mosquito Alert app. This type of information is of a different nature from the field sampling and still needs expert interpretation, but offers significantly broader and faster geographical coverage. Two decades have passed since the first detection of the Asian tiger mosquito in Spain in 2004, and following the Mosquito Alert creation in 2014, citizen science has contributed with one-third of the discoveries of this species at the municipal level in Spain. We discuss integration of both approaches in order to harness their respective strengths while compensating their drawbacks, to develop faster and more powerful early warning surveillance systems, and also returning value to the society by empowering citizenship.

## 1. Introduction

### Context

The global expansion of certain mosquito species (Diptera: Culicidae) results from a combination of their adaptive capacity and multiple human-related factors, including increased international mobility, trade, and climate change [1]. While the global issue of invasive species primarily impacts biodiversity and economy, it also has significant public health implications in the case of invasive mosquitoes that act as vectors of animal and human diseases. Besides of the species dispersal, the aforementioned factors also contribute to the redistribution of associated pathogens—particularly through increased mobility of infected individuals to and from endemic areas. This heightens the likelihood of pathogen-vector overlap, thereby increasing the risk of local disease transmission, of which there are already precedents in Europe and specifically in Spain [2]. Even in the absence of transmission, the aggressiveness, anthropophily, and domestic presence of some mosquito species constitute a nuisance that diminishes people’s quality of life, a relevant component of public health.

Despite its growing visibility in the public sphere, this issue has deep historical roots. During the 17^th^ century, the infamous slave trade from West Africa to the Americas facilitated the transport of *Aedes aegypti*, a mosquito species with high vector competence for human viruses such as yellow fever (YFV) and dengue (DENV) which were transported in parallel [3]. Later, during what may be considered as an early phase of globalization from 17^th^ to 20^th^ centuries, the intense commercial exchange between the Americas and Europe resulted in frequent arrivals to Spain of infested ships, along with crews infected by these viruses. Disembarkation events frequently triggered port outbreaks that sometimes lasted for years and reached distant populations—for instance, the series of yellow fever outbreaks between 1800 and 1803 that caused over 60,000 deaths in Cádiz, Jerez de la Frontera, and Seville [3]. It is estimated that more than 300,000 deaths in Spain during the first half of the 19^th^ century were attributable to this cause [4]. As *Ae. aegypti* is one of the most globally significant vector species, its ecology has been extensively studied under a control point of view [5]. Nevertheless, significant knowledge gaps remain regarding the factors that led to its disappearance from most of Europe during the first half of the 20^th^ century, probably due to a combination of technological, social, environmental and health-related changes which require further investigation, given the striking disparity between its estimated potential distribution and its currently limited presence in Europe [6]. The species’ persistence in Eastern Europe, its reintroduction in Madeira in 2005 [7] (followed by a dengue outbreak a few years later [8]) and its recent detections in Cyprus [9] and the Canary Islands [10] suggest that this period of apparent absence from Europe may be temporary. For this reason, *Ae. aegypti* currently represents a top priority for entomological surveillance of invasive vectors.

The Asian tiger mosquito (*Aedes albopictus*) which is the invasive mosquito species widely established in the Mediterranean, is undergoing a process of global expansion that began in Europe in Albania in 1979 [11], and in Spain from 2004 onwards [12]. This species is a competent vector for dengue, chikungunya (CHKV), and Zika (ZIKV) viruses among other pathogens, and poses a major public health risk due to its adaptive capacity, including hibernation, and proven invasive potential [13]. At present, it constitutes the primary risk factor for these three exotic arboviral diseases in Spain.

The third invasive mosquito species identified in Spain is *Aedes japonicus*, first discovered in Asturias in 2018 through citizen science [14] and later detected in extensive areas across northern Spain [15]. This species had not initially been the subject of targeted surveillance, as its nearest known populations were located over a thousand kilometers away in North-Eastern France. Its vector potential is considered relatively low. Laboratory experiments have shown it is competent for West Nile virus (WNV) [16] though several more efficient native vectors already exist [17]. Its dispersal potential in the Iberian Peninsula is also thought to be limited due to its preference for environmental conditions typical of continental climates [18].

Finally, it is worth noting the existence of *Aedes koreicus*, a species closely related to the former, currently limited to central European regions. Although it has not yet been detected in Spain and is considered to have low vectorial capacity for the mentioned pathogens [19], it warrants attention due to the precedent set by the demonstrated invasive potential of *Ae. japonicus*.

### The administrative framework of surveillance

The public health risk posed by invasive vectors needs the implementation of entomological surveillance protocols to enable early detection in yet uncolonized territories and—to the extent that species are already established—to provide data to support vector control and public health interventions.

Administrative responsibility for mosquito-related matters typically falls under the public health umbrella, although it is often shared with other areas of government due to the intersection with animal health, vector management, and even biodiversity protection in the case of *Ae. albopictus*, which is listed in the Invasive Species Catalog established under the Spanish Law 42/2007 on Natural Heritage and Biodiversity Protection [20]. This highlights the need for interdepartmental collaboration as an integral component of cross-sectoral approaches within the One Health framework [21].

Being a highly decentralized country, the executive duties in Spain are also distributed across levels of administration, from municipalities to the State. Entomological surveillance is not legally mandated and is the competence of 19 regional autonomous entities -namely 17 Autonomous Communities and 2 Autonomous Cities (all collectively referred thereafter as ACs). Most of these have developed management plans for vectors and diseases with the support of the healthcare sector, academic institutions, public mosquito control services and other stakeholders [22–28], as awareness of its importance developed in parallel with the progressive colonization of new regions by the tiger mosquito.

Since 2008, the Spanish Ministry of Health has played a major role in this process carrying out surveillance activities in ports and airports [29] in accordance with the International Health Regulations (IHR) [30] which mandate monitoring at points of entry. Through the Coordination Centre for Health Alerts and Emergencies (CCAES) and the Sub-Directorate for Environmental Health, the Ministry also plays an informational and coordinating role among the ACs, conducting annual surveys to compile data on surveillance, establishment, and control of vectors at the municipal level, and publishing an annual report since 2016 [31]. Finally, the Ministry also coordinates the National Plan for the Prevention, Surveillance and Control of Vector-Borne Diseases (PNPVC) [32].

Academic research has contributed review studies about distribution [33,34], and as of 2023, Spain participates through Vectornet-ES [35] to the European VectorNet network [36], coordinated by the European Centre for Disease Prevention and Control (ECDC) and the European Food Safety Authority (EFSA). These institutions manage the European vector distribution databases connected to maps [37] and promote advisory, preparedness, and response collaborations for vector-borne diseases.

### Field surveillance

Field surveillance methodologies generally follow ECDC guidelines [38] including larval sampling, ovitraps, and devices for capturing adult mosquitoes. However, these techniques are not standardized to protocols, despite some intense efforts like the pioneering European initiative AIMSurv [39]. Field-based surveillance provides reliable physical evidence but lacks speed, is constrained by scale, and is resource-intensive, requiring trained personnel for both sample collection and laboratory identification. Therefore, careful selection of monitoring areas is critically based on cost-benefit considerations and risk hypotheses, particularly regarding points of entry. A common strategy is to monitor municipalities adjacent to already affected areas, assuming a local dispersal by autonomous flight and passive transport in vehicles [40]. As short-range dispersal is rapid and nearly unavoidable, this approach yields effective results at the regional level. However, it does not fully account for the broader process, as the stratified dispersal pattern of these species spans a wide range of spatial scales [41] with local contagious spread being just one of them. In addition, long-distance dispersal events are associated with commercial transport, are not continuous in time, involve large numbers of immature stages at once, and transcend the limited jurisdictional boundaries in a unified market. Hence, the unpredictability of these dispersal directions poses a challenge to surveillance by field sampling which is often limited to target entry points and areas associated to transportation networks, such as bus terminals and highway service stations [42]. Despite its inherent limitations in cost and scale, conventional entomological sampling remains a unique and indispensable tool, providing physical evidence classified with a high degree of certainty by trained specialists.

### Citizen science surveillance

Ideally, surveillance systems should be capable of producing big data in real time, and with a level of scalability that mirrors the rapid pace of mosquito dispersal. These are key features of citizen science, made increasingly feasible by the widespread use of powerful mobile devices with integrated cameras. The involvement of thousands of individuals generating large volumes of data from any location overcomes limitations related to scale, cost, and time, and is particularly effective for early warning and for mapping remote invasion fronts [43].

The Mosquito Alert citizen science project (https://www.mosquitoalert.com), launched in 2014 in Spain by public research institutions (initially under the name Tigatrapp), offers a digital platform with a mobile app that enables users to submit photographic reports of mosquitoes. Initially focused on the tiger mosquito (*Ae. albopictus*), the project later expanded its scope to include the yellow fever mosquito (Ae. aegypti), the Japanese bush mosquito (*Ae. japonicus*), the Korean mosquito (*Ae. koreicus*), and the common house mosquito (*Culex pipiens s*.*l*.), the latter not being invasive but of interest due to its role in transmitting WNV among other diseases of interest under the One Health framework. The project also escalated to 23 European states in the framework of AIMCost [44] and VEO projects [45], is open-source (https://github.com/Mosquito-Alert), all users are completely anonymous and provide their data to be available under a Creative Commons Zero - CC0 1.0 license. Data can be downloaded from a portal (https://labs.mosquitoalert.com/metadata_public_portal/) but are also available in the project’s interactive web map (https://map.mosquitoalert.com), refreshed on a daily basis. Photographic reports of adult mosquitoes are classified in the EntoLab, a web platform that combines artificial intelligence (AI) analysis with expert validation. For each incoming report the AI provides an initial classification, which is sent to the user’s device, followed later by the evaluation of the three experts specialized in Culicids and selected on a regional basis from a global pool of 137 volunteers: 25 in Spain, 83 in other European countries and 29 in local implementations worldwide.

Due to its extensive coverage and scalability, citizen participation is especially suited for early warning systems, though its utility goes beyond that. The opportunistic sampling associated with citizen reports can be combined to entomological field data, sociological parameters and environmental information to build vector dispersal and risk models, valuable for both vector management and epidemiological risk assessment (see for example https://labs.mosquitoalert.com/MosquitoAlertBCN/). The system includes a procedure for estimating the sampling effort of participants over time and per geographic unit that allows to cancel out opportunistic sampling in model predictions of citizen exposure to mosquitoes, and derived vector suitability maps. The sampling effort modelling was not used in the present study, which aims only to reflect citizen-sourced discoveries of invasive species in new territory.

Given these capabilities, citizen science was, for the first time in Europe, formally integrated into the Spanish PNPVC in 2023 as a method for near-real-time monitoring, assessment, and early warning. The quality of citizen-generated data, however, is not exempt from uncertainties, such as the need to interpret digital evidence and the potential mislocation of reports and images. Under a big data approach, these inaccuracies are expected to be compensated by the large volume of information, which minimizes the potential bias of individual records, or by field surveillance verification. Another challenge is maintaining citizen engagement. Effective surveillance requires long-term participation, with consistent reporting throughout each mosquito season, year after year. Achieving this level of commitment demands well-designed communication strategies—both short- and long-term—as well as sustained efforts to keep participants motivated over time. In this context, the Mosquito Alert platform includes communication tools to participants, enabling the dissemination of notifications to individual devices, or broadly across users in specific geographic areas. Together with the platform’s web publications and social media presence, this completes a comprehensive cycle of engagement, information-sharing, and citizen empowerment regarding vector-related challenges.

### Aiming to build combined listings and protocols

Dispersion of information across sources has hindered the updating and consolidation of a municipal reference list of invasive mosquito detections, which is an essential tool for the community. The last academic review on distribution was published in 2015 [33] and from 2016 onwards the Ministry of Health is publishing annual reports with lists of municipalities compiled through questionnaires completed by AC sources. Citizen Science is contributing from 2014 to this task, although there are no centralized repositories integrating both data sources which would allow for a reliable, accessible and complete dataset.

Therefore, the first aim of this work is to compile the distribution list of invasive mosquito species, combining data from both sources: field sampling (2004–2024) and citizen science (2014–2024). Considering the ten-year period of overlapping operation of both surveillance strategies, we also compare these two detection approaches in terms of speed and spatial coverage. Faster detection facilitates rapid response and potential eradication, and enables more accurate anticipation of epidemiological risk; geographical range is particularly relevant to risk assessment and for improving vector surveillance management strategies. Finally, we discuss the potential for future integration of the two surveillance strategies in order to compensate their differences while complementing their strengths.

## 2. Materials and Methods

The study period spans 20 years from the detection in 2004 of Ae. albopictus (the first invasive mosquito species in Spain) to the present. The compiled dataset records the year of the first known detection per municipality and species, specifying whether the detection was made via digital citizen reporting, field sampling, or both. The global temporal resolution has been limited to the calendar year because of the absence of precise temporal information within the published field data. However, for citizen science reports, the exact day is available and can be checked in the public Mosquito Alert map.

### Field sampling

All available data for field sampling were compiled, including the authors’ own datasets, institutional documents, data from academic and administrative territorial stakeholders, regional lists published online and scientific literature. Most published listings specified only the detection year and the location, with no mention of specific date, capture method nor sampled lifestage. Author’s own data were obtained by ovitrapping and raising the eggs to adult stage, larval sampling, genetic analysis on eggs, or direct adult examination. Species identification was performed via appropriate taxonomic keys [46]. Necessary reconciliations were made in case of discrepancies, giving priority to peer-reviewed scientific publications and the authors’ own data. While unlikely, information gaps in field information may exist for the 2022–2024 period, particularly if certain results have not yet been published. The institutions contributing data to this work are listed in Table 1.

**Table 1.** Administrative bodies having directly supplied data listings to the authors for this study.

|  |  |
| --- | --- |
| Catalonia | * Departament de Salut |
| Generalitat de Catalunya | * Departament d'Interior i Seguretat Pública |
|  | * Departament de Territori, Habitatge i Transició Ecològica |
| Andalusia | * Servicio de Salud Ambiental, Dirección General de Salud Pública y Ordenación Farmacéutica, Consejería de Salud y Consumo |
| Junta de Andalucía |  |
| Extremadura | * Dirección General de Salud Pública, Servicio Extremeño de Salud |
| Junta de Extremadura |  |
| Galicia | * Dirección General de Salud Pública, Consellería de Sanidade |
| Xunta de Galicia |  |
| La Rioja | * Servicio de Salud Ambiental y Nutrición, Dirección General de Salud Pública, Consumo y Cuidados, Consejería de Salud y Políticas Sociales |
| Gobierno de La Rioja |  |
| Navarre | * Instituto de Salud Pública y Laboral de Navarra (ISPLN) |
| Comunidad Foral de Navarra |  |
| Aragon | * Servicio de Seguridad Alimentaria y Sanidad Ambiental, Dirección General de Salud Pública |
| Gobierno de Aragón |  |
| Basque Country | * Departamento de Sanidad Animal de NEIKER - Instituto Vasco de investigación y desarrollo agrario |
| Eusko Jaurlaritza | * Dirección de Salud Pública y Adicciones |
| Madrid | * Área de vigilancia de riesgos ambientales en salud, Consejería de Sanidad |
| Comunidad de Madrid |  |
| Murcia<br>Comunidad Autónoma de la<br>Región de Murcia | * Servicio de Sanidad Ambiental, Dirección General de Salud Pública y Adicciones |
| Ministry of Health<br>Ministerio de Sanidad | * Centro de Coordinación de Alertas y Emergencias Sanitarias (CCAES)<br>* Servicio de Riesgos Ambientales, subdirección General de Sanidad Ambiental |

### Citizen science

All citizen reports geolocated within Spanish municipalities received between the launch of Mosquito Alert on June 6, 2014, and December 31, 2024 were compiled. This dataset comprises a total of 110,939 reports across three categories (adult mosquitoes, breeding sites, or biting activity), submitted by 33,183 participants. Only reports showing an adult mosquito (of either sex) were considered. The data are publicly accessible and can be downloaded via the online map, except for those reports removed due to private or inappropriate content.

Validation of each report by three expert entomologists relied on a set of three species-specific morphological criteria not related to sex, whose combination allowed the classification of a picture as either “probable” or “confirmed” for *Ae. albopictus, Ae. aegypti*, or *Ae. japonicus*. Additional categories were also available, such as “other species” (whether belonging to *Culicidae* or not), or null answers. Consensus among the three evaluations was automatically calculated by the system by weighted integration of individual outputs, after which a supervisor reviewed all results for consistency. Consequently, a municipality was considered citizen science-positive for a given year and species when at least one report from its territory was classified as “probable” or “confirmed” for that species.

Only the year of the first detection is included in the listing, with subsequent detections omitted, as the aim was to identify the temporal onset of invasive species rather than to assess their establishment. Therefore, no quantitative parameters regarding sustained presence or inter-report intervals were analyzed. Due to the intrinsic uncertainty associated with citizen-generated data, cases in which a municipality’s positive status was based on a single report were identified and tagged for evaluation.

### Integration into the database

Each entry in the dataset corresponds to a municipality at the LAU2 level (formerly NUTS level 5), identified by its administrative code, along with its autonomous community and province, population (2024) and municipal surface area in hectares. The descriptive section includes nine data columns, three for each species. For each species, the first column records the year of detection via field sampling, if available; the second column contains the source of that information—either a bibliographic reference or, in the case of unpublished material, the designation “own data”; the third column lists the year of detection via citizen science, if applicable.

As a result, five possible categories could be defined per species for each municipality: (1) positive by field sampling only; (2) positive by citizen science only; (3) positive by both strategies in the same year; (4) positive by both strategies in different years; and (5) no detection. The strategy attributed to the first detection corresponds to the earliest year between columns one and three; if both years are equal, the municipality is classified as having simultaneous detection from both sources. However, in simultaneous cases where the chronological sequence of events was well established, the original surveillance strategy was retained as the primary source of the event.

### Geographic tools

The Spanish list of municipalities is based on the official database of the National Statistics Institute, updated as of January 1, 2024 [47] and comprising a total of 8,132 municipalities. In addition, there are 86 non-municipal territorial entities—such as *mancomunidades, ledanías, facerías, parzonerías*, common lands, islets, and other unique or cooperative territorial units—which, although not municipalities, are also assigned identification INSPIRE codes (ES.IGN.BBDAE), so that they are included in geographic representations for consistency with the used cartography, obtained from the National Geographic Institute [48] used in QGIS 3.34.10 at the ETRS89 reference system, under a stereographic projection ESRI:54026.

### Human population density and the spread of *Aedes albopictus*

To evaluate the progression of the spread in relation to local demographics characteristics, we analyzed the population density in municipalities where Ae. albopictus was newly detected each year. Statistical significance was assessed using a province-stratified permutation test with 499 iterations and equal sample size: for each year, the set of newly detected municipalities was compared to a spatially matched null distribution generated by randomly sampling an equal number of non-detected municipalities, preserving the same relative representation of provinces. The test was not performed for 2004, the year of first detection, as only two municipalities were affected. Surface area and population data were obtained from the Geographic Nomenclature of Municipalities and Population Entities, published by the National Geographic Institute [49] under CC-BY 4.0 license.

### Detection distance for *Aedes albopictus* by strategy

Geographic reference points were determined for each municipality by computing the geometric centroids of their polygon using the *Centroids* plugin in QGIS. For each newly positive municipality for *Ae. albopictus*, and for both surveillance strategies (field sampling and citizen science), the distance in kilometers was calculated to the nearest municipality that had been positive in the previous year. Distances were computed in R using the *distVincenty* function from the Geosphere package [50], based on Vincenty’s spherical method [51]. Specific adjustments were made in exceptional cases: for the five municipalities in Galicia detected in 2023, the distance assigned was relative to Penafiel (Portugal), where an established *Ae. albopictus* population had been known since 2017 [52], (centroid coordinates: latitude = 41.20683, longitude = –8.28486). North African and insular municipalities were retained in the analysis, excluding the two in the Canary Islands, where all *Ae. albopictus* detected populations were considered eradicated. We also excluded all municipalities with simultaneous detections in the same year, the two initial municipalities from 2004 (due to the absence of prior reference points), and the municipality of Orihuela (Alicante), detected in 2005 at over 560 kilometers from the initial records and being suspected of an independent introduction [53].

A two-sample Student’s t-test was applied to the mean distances, using the surveillance strategy (field sampling vs. citizen science) as the grouping factor. Additionally, a two-way ANOVA was performed in R with sampling year and strategy as factors and post-hoc Tukey group comparisons. The analysis was repeated including the municipalities detected via field sampling between 2004 and 2013, and it was also verified that the applied exclusions did not alter the statistical significance of the results.

## 3. Results

The number of detections of at least one of the three invasive mosquito species by any of the two surveillance strategies sums a total of 1,813 municipalities in Spain, representing 22.3% of the country’s 8,132 municipalities. A summary of the municipality counts by species is presented in Table 2, and their geographic distribution is shown in Figure 1. The final municipal dataset is available at https://doi.org/10.5281/zenodo.15869762.

**Table 2.** *Number of municipalities with detections of any of the three* Aedes *mosquito invasive species. Four of the listed detections are currently considered eradicated, all in the Canary Islands: two of Ae. albopictus and two out of three detections of Ae. aegypti.*

| *Species* | *Municipalities* |
| --- | --- |
| *Aedes albopictus* | 1,699 |
| *Aedes japonicus* | 43 |
| *Ae. albopictus* and *Ae. japonicus* | 68 |
| *Aedes aegypti* | 2 |
| *Ae. albopictus* and *Ae. aegypti* | 1 |
| **TOTAL** | **1,813** |

**Figure 1.**
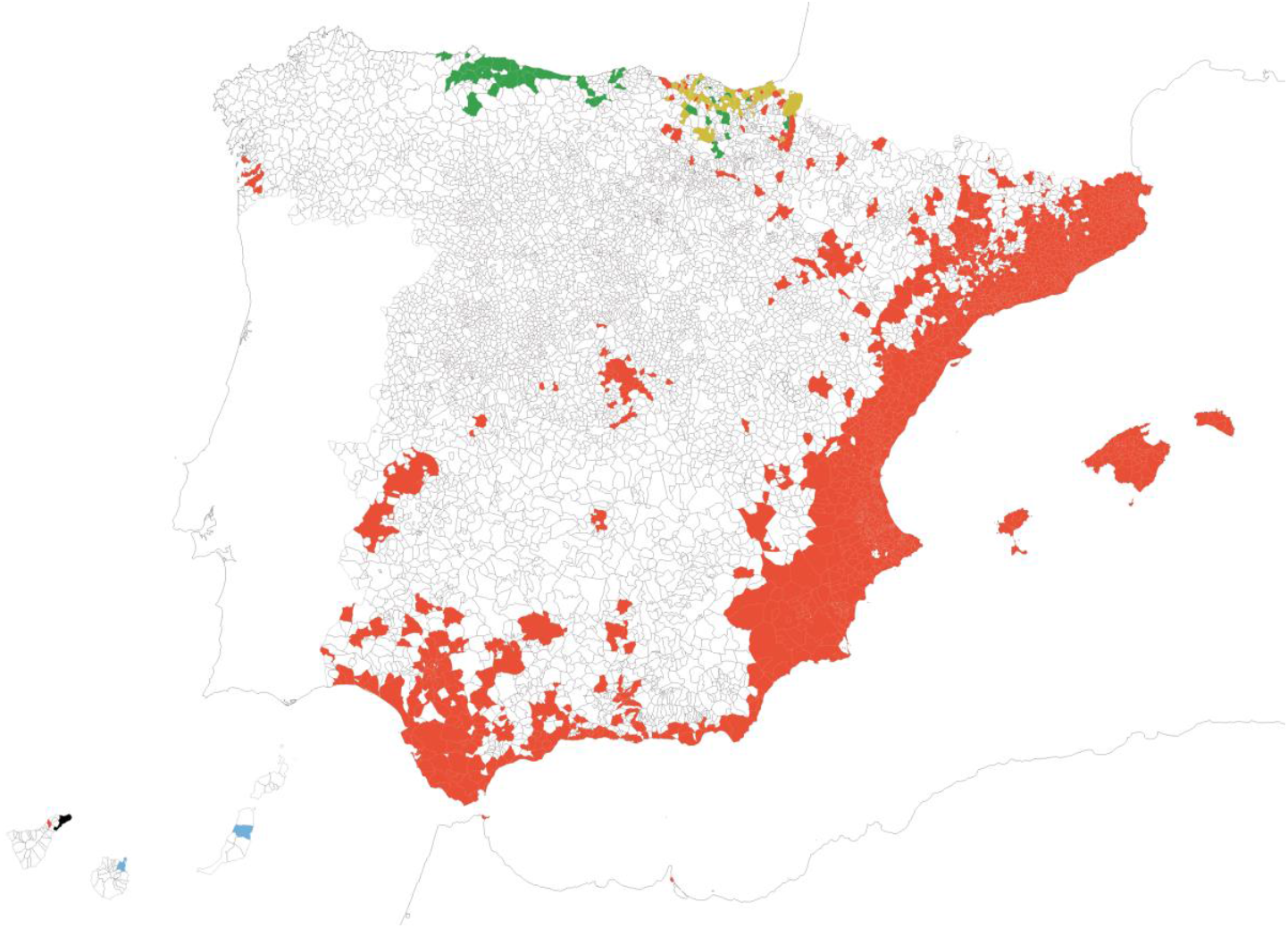
Map of Spanish municipalities with reported detections during the period 2004–2024 of Ae. albopictus alone (N=1,699, red), Ae. japonicus alone (N=43, green), overlapping detections of Ae. albopictus/Ae. japonicus (N=68, ochre), Ae. aegypti alone (N=2, blue), and overlapping Ae.albopictus/Ae. aegypti (N=1, black). Map reference: BDLJE CC-BY 4.0, National Geographic Institute – Canary Islands displaced from their actual position.

The following section presents species data, with particular emphasis on *Ae. albopictus*, as it is by far the most prevalent and relevant invasive species at present due to its territorial, health, and social impact.

### Aedes japonicus

The first record of *Ae. japonicus* in Spain was obtained through Mosquito Alert, based on an alert issued in Asturias in June 2018 [14]. Following its discovery—and after a mass notification sent to all users in northern Spain—Mosquito Alert contributed to the subsequent delineation of its distribution [15]. To date, the species has been reported in 111 municipalities within the ACs of Asturias, Cantabria, the Basque Country, and Navarre, in an effective collaboration between citizen scientists and field entomologists. These municipalities represent 1.4% of the total number in Spain, covering an area of 6,588 square kilometers (1.3% of the national surface) and a combined population of 3,097,093 inhabitants (6.4% of the national population). Discovery at each municipality was attributed to field sampling in 66.7% of cases, Mosquito Alert in 24.3%, and simultaneous detection through both sources in 9%. In 68 of the 111 municipalities, the species co-occurs with *Ae. albopictus*.

### Aedes aegypti

In recent years the yellow fever mosquito has been detected in the Canary Islands in 3 main events: by 2017, it was identified through field sampling in Puerto del Rosario, Fuerteventura [54] from where it was declared eradicated in 2019 [55]. Regarding Mosquito Alert, two reports were submitted: one in 2022 from Santa Cruz de Tenerife (also considered eradicated), and another in December 2023 from the Piletas neighbourhood of Las Palmas de Gran Canaria, which was simultaneous to a field detection [56]. Special considerations are warranted for this autonomous community, given its heightened exposure to repeated introductions of invasive *Aedes* species via maritime routes from Madeira and Africa. While it maintains a robust surveillance and control system, meaningful citizen science resources have not yet been actively promoted within the region.

### Aedes albopictus

*Ae. albopictus* has been detected in 1,768 municipalities, accounting for 21.7% of the total number of municipalities in Spain. These cover a combined surface of 108,626 square kilometers, totalizing a 21.5% of the national territory. However, the resident population affected totals 31,789,424 people, representing a significantly higher share of 66.2% of the total population. Table 3 presents the total number of Ae. *albopictus*-positive municipalities per autonomous community, alongside basic data on surface area and population, as well as national totals. At the regional level, Asturias appears to be the only AC that remains free of *Ae. albopictus* to date, together to Canary Islands if considering eradications.

**Table 3.**
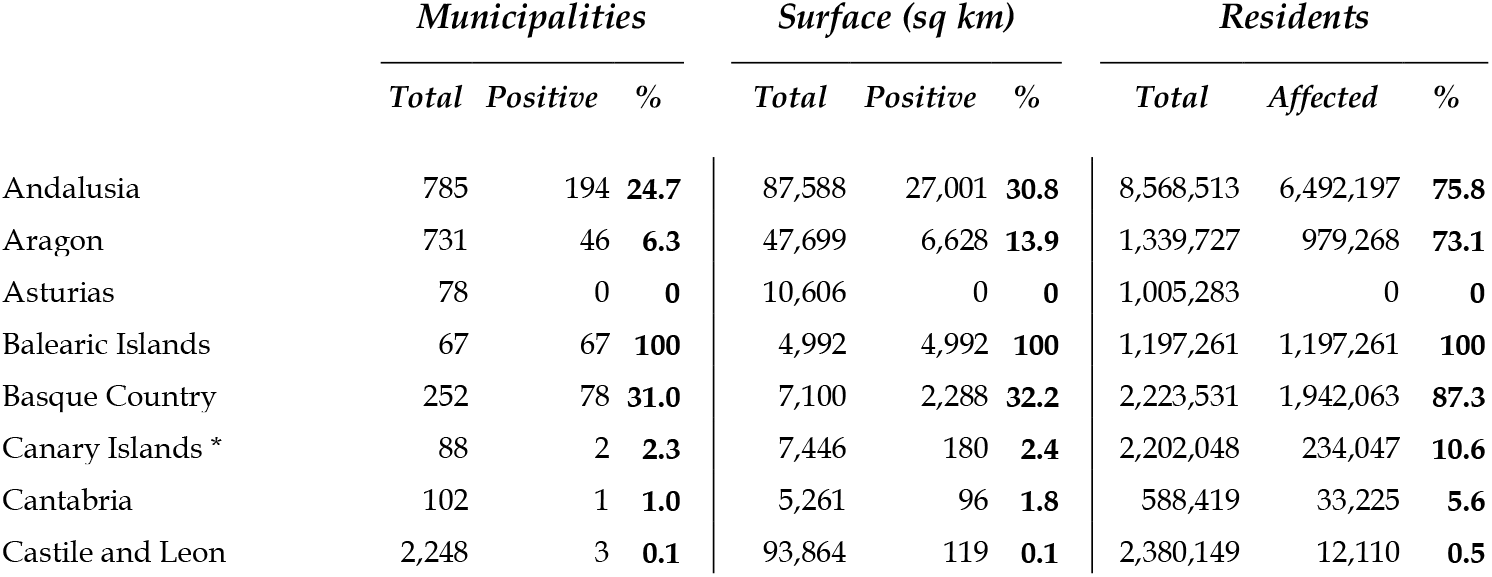
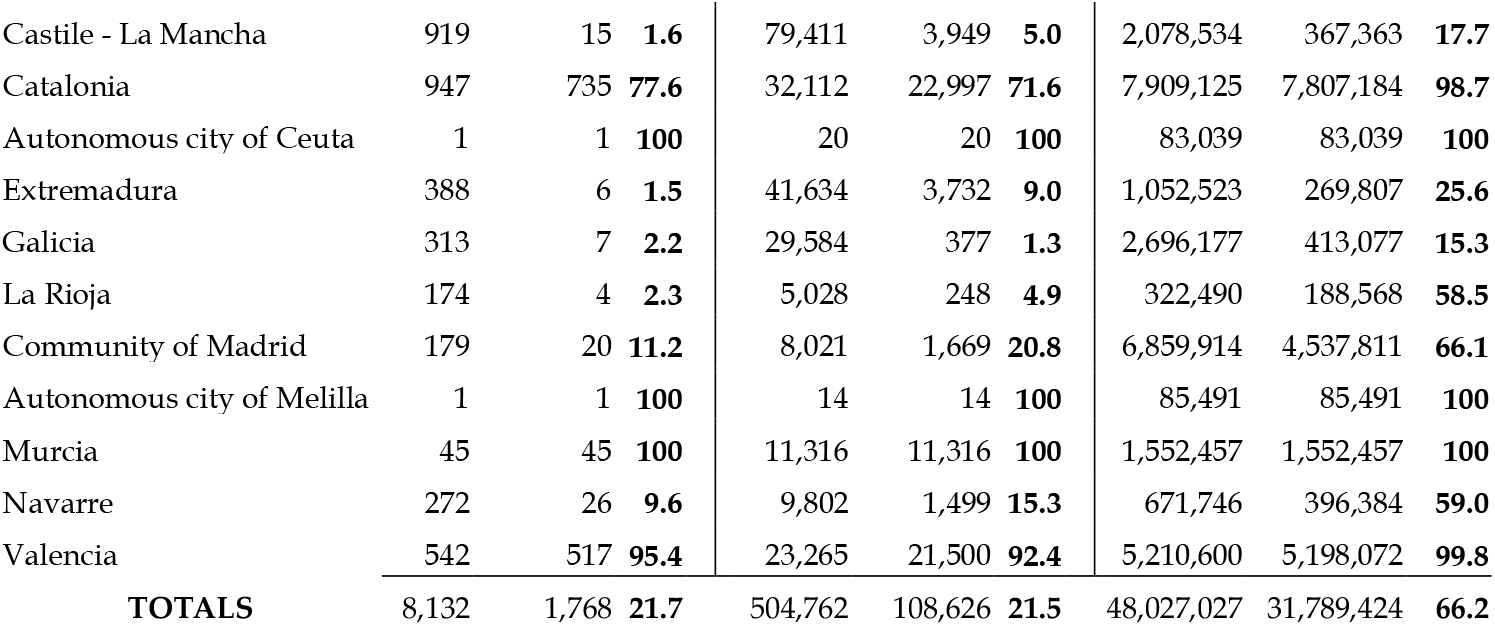
Municipalities, surface areas, and populations affected by Ae. albopictus relative to the totals, broken down by autonomous community. * Assuming eradications, the real affected proportions in Canary Islands should all be zero for Ae. albopictus.

|  | *Municipalities* |  |  | *Surface (sq km)* |  |  | *Residents* |  |  |
| --- | --- | --- | --- | --- | --- | --- | --- | --- | --- |
|  | *Total* | *Positive* | *%* | *Total* | *Positive* | *%* | *Total* | *Affected* | *%* |
| Andalusia | 785 | 194 | **24.7** | 87,588 | 27,001 | **30.8** | 8,568,513 | 6,492,197 | **75.8** |
| Aragon | 731 | 46 | **6.3** | 47,699 | 6,628 | **13.9** | 1,339,727 | 979,268 | **73.1** |
| Asturias | 78 | 0 | **0** | 10,606 | 0 | **0** | 1,005,283 | 0 | **0** |
| Balearic Islands | 67 | 67 | **100** | 4,992 | 4,992 | **100** | 1,197,261 | 1,197,261 | **100** |
| Basque Country | 252 | 78 | **31.0** | 7,100 | 2,288 | **32.2** | 2,223,531 | 1,942,063 | **87.3** |
| Canary Islands * | 88 | 2 | **2.3** | 7,446 | 180 | **2.4** | 2,202,048 | 234,047 | **10.6** |
| Cantabria | 102 | 1 | **1.0** | 5,261 | 96 | **1.8** | 588,419 | 33,225 | **5.6** |
| Castile and Leon | 2,248 | 3 | **0.1** | 93,864 | 119 | **0.1** | 2,380,149 | 12,110 | **0.5** |
| Castile - La Mancha | 919 | 15 | 1.6 | 79,411 | 3,949 | 5.0 | 2,078,534 | 367,363 | 17.7 |
| Catalonia | 947 | 735 | 77.6 | 32,112 | 22,997 | 71.6 | 7,909,125 | 7,807,184 | 98.7 |
| Autonomous city of Ceuta | 1 | 1 | 100 | 20 | 20 | 100 | 83,039 | 83,039 | 100 |
| Extremadura | 388 | 6 | 1.5 | 41,634 | 3,732 | 9.0 | 1,052,523 | 269,807 | 25.6 |
| Galicia | 313 | 7 | 2.2 | 29,584 | 377 | 1.3 | 2,696,177 | 413,077 | 15.3 |
| La Rioja | 174 | 4 | 2.3 | 5,028 | 248 | 4.9 | 322,490 | 188,568 | 58.5 |
| Community of Madrid | 179 | 20 | 11.2 | 8,021 | 1,669 | 20.8 | 6,859,914 | 4,537,811 | 66.1 |
| Autonomous city of Melilla | 1 | 1 | 100 | 14 | 14 | 100 | 85,491 | 85,491 | 100 |
| Murcia | 45 | 45 | 100 | 11,316 | 11,316 | 100 | 1,552,457 | 1,552,457 | 100 |
| Navarre | 272 | 26 | 9.6 | 9,802 | 1,499 | 15.3 | 671,746 | 396,384 | 59.0 |
| Valencia | 542 | 517 | 95.4 | 23,265 | 21,500 | 92.4 | 5,210,600 | 5,198,072 | 99.8 |
| **TOTALS** | **8,132** | **1,768** | **21.7** | **504,762** | **108,626** | **21.5** | **48,027,027** | **31,789,424** | **66.2** |

Figure 2 presents the temporal evolution over the study period regarding affected surface area, number of inhabitants, and municipalities, in this case in accumulated form and also as annual new additions. Along the first 9 years (2004-2013) based only on field sampling, the mean number of municipalities detected per year was 40.5, mostly mirroring the field-based monitoring in Catalonia, and adjusting well to a linear trend (R^2^=0.9572). From 2014 to 2024 the inclusion of citizen science and enhancement of surveillance programs raised this figure to 123.9 municipalities per year, although under a higher inter-year variability departing to the linear fit (R^2^=0.130) with marked lows in 2017 and 2022. Figure 3 displays the geographic spread of *Ae. albopictus* over time.

**Figure 2.**
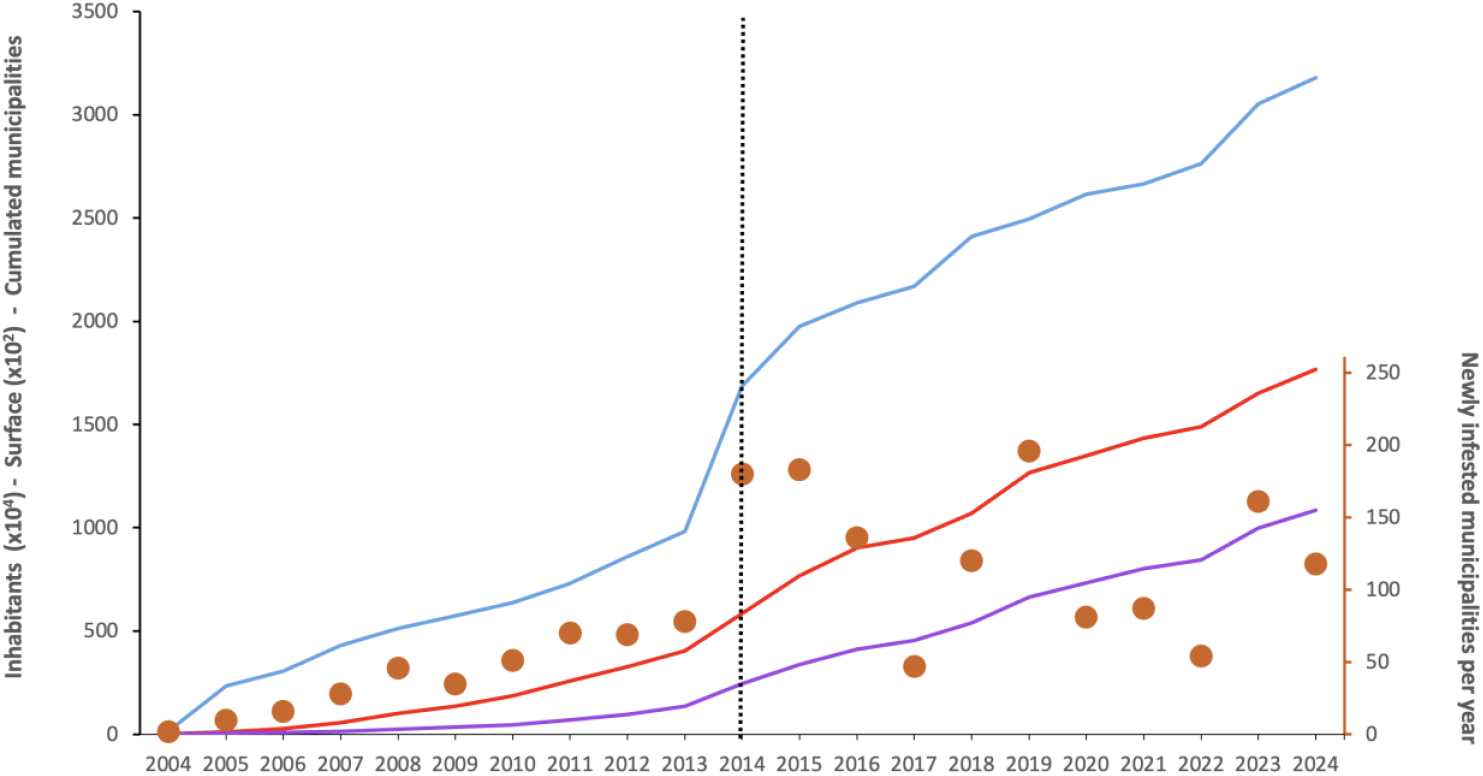
Twenty years of Ae. albopictus in Spain: cumulative number of affected inhabitants (blue line, transformed by a factor of 1×10^−4^ for ease of visualization), cumulative occupied area in square kilometers (purple line, transformed by a factor of 1x10^−2^), and cumulative number of municipalities (red line). The dot series refers to the secondary vertical axis (right) and shows the number of newly infested municipalities per year. The first detection of invasive mosquito species was in 2004 and the start of Mosquito Alert activity in 2014 is indicated by the vertical dashed line.

**Figure 3.**
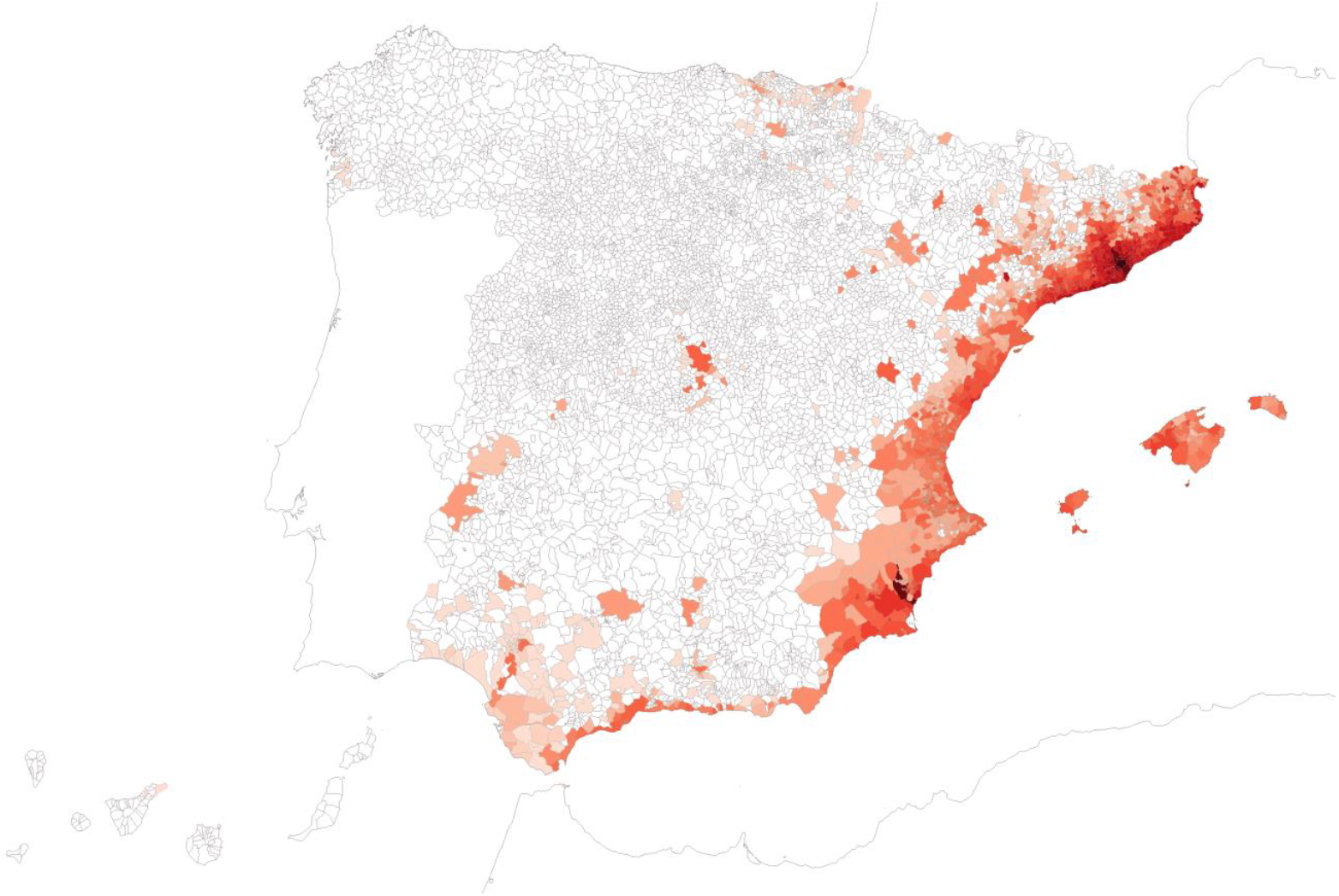
Detections of Ae. albopictus by any strategy, broken down by year of first detection. Darker shades indicate earlier detections, starting 2004 up to 2024. Map reference: BDLJE CC-BY 4.0, National Geographic Institute – Canary Islands displaced from their actual position.

### Human population density and the spread of *Aedes albopictus*

A Student’s T-test revealed no significant differences in municipal population densities between surveillance strategies (P>0.4). Therefore, the analysis was conducted on the pooled dataset as shown in Figure 4, where each boxplot summarizes the distribution of population densities in municipalities where the species was first detected in a given year. The horizontal line inside each box represents the median, boxes indicate the interquartile range (IQR), and whiskers extend to 1.5× IQR. Jittered dots represent individual municipalities, providing a view of the underlying data distribution. As the species colonizes more municipalities over time, the average population density of newly invaded areas gradually converges toward the average municipality density in Spain (182.49 inhab/ha, indicated by the dashed horizontal line). Whereas considerable variation remains across years, only in 2013 and 2021 the invaded municipalities did not exhibit significantly higher population densities compared to their matched, non-invaded counterparts.

**Figure 4.**
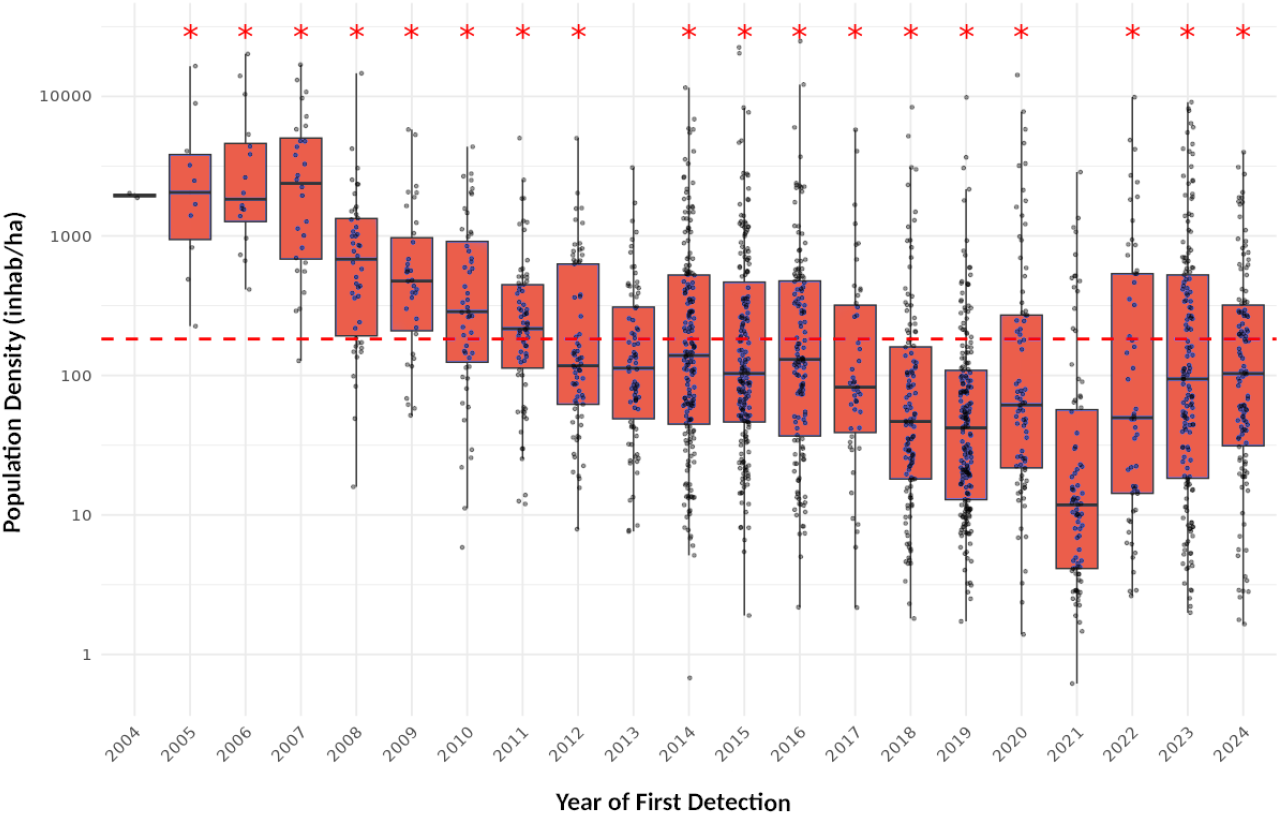
Population density (log_10_ transformed) of Spanish municipalities colonized by Ae. albopictus, by year of detection. The red dashed horizontal line marks the overall mean population density across all municipalities, serving as a reference point for urbanization level. Asterisks indicate the level of statistical significance for the test, showing whether the detected municipalities had significantly higher population density than expected by chance (P < 0.05)

### Detections of *Aedes albopictus* by surveillance strategy

The substantial surveillance efforts carried out by academic institutions and government agencies are evident from the results since 2004, with authoritative field-based detections in 1,335 municipalities by 2024, though some of them were poorly attributed to years in some reports. Probably due to decentralized management, access to results is relatively limited as data are released with notable delays, often exceeding one year.

In parallel, Mosquito Alert contributed 335 additional detections during its operational period from 2014 to 2024. During this shared timeframe, 98 more municipalities were identified as positive by both strategies in the same year, posing challenges for interpretation. These results are summarized in Table 4, represented in the map in Figure 5, and plotted by year in Figure 6.

**Table 4.**
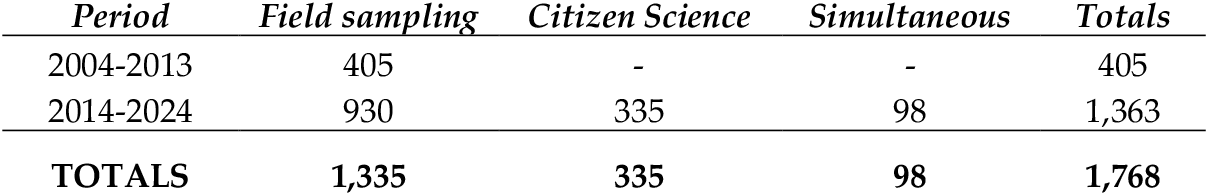
Number of first municipal detections of Ae. albopictus, broken down by surveillance strategy and period.

**Figure 5.**
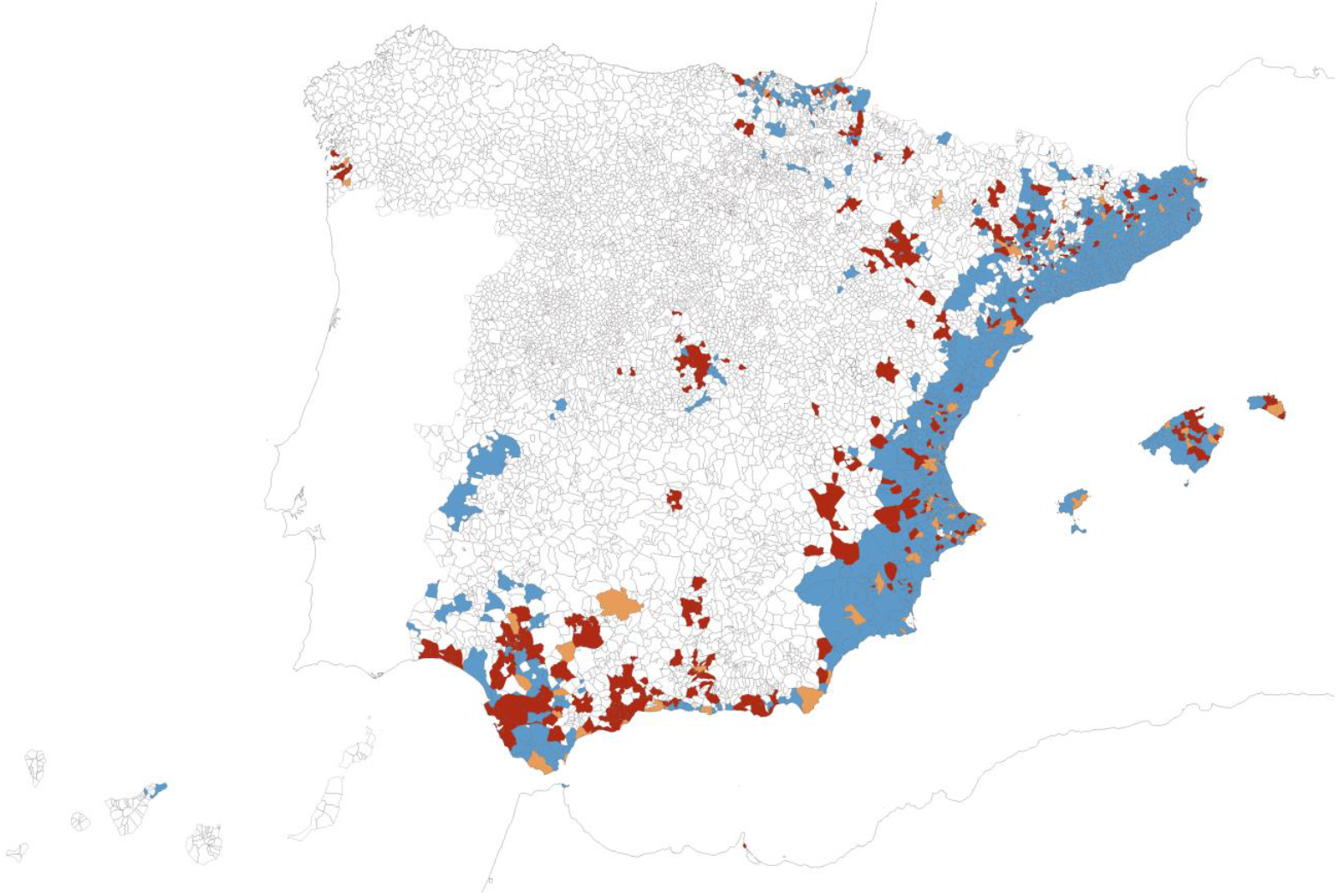
Records of Ae. albopictus, categorized by surveillance strategy (blue: field sampling 2004-2024, red: citizen science 2014–2024, ochre: simultaneous reports from both sources). Map reference: BDLJE CC-BY 4.0, National Geographic Institute – Canary Islands moved from their actual position.

**Figure 6.**
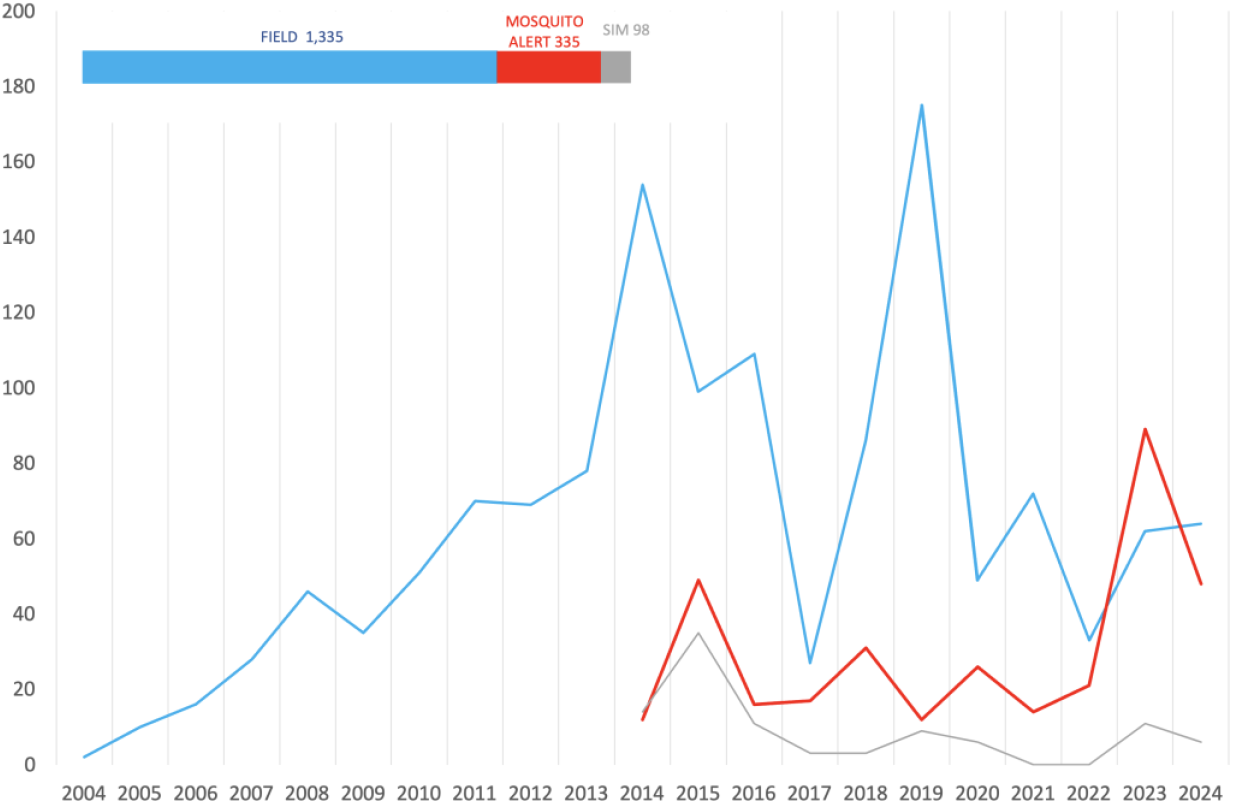
Number of municipal detections of Ae. albopictus per year and strategy (2004-2024). Blue: field sampling, red: Mosquito Alert, grey: simultaneous discoveries. Inset: the bar sections represent the proportion of each class along the whole study period.

Along the overlapping period, Mosquito Alert was responsible for one quarter of the 1,363 total detections (24.6%), and when simultaneous detections are included, the platform was involved in 31.8% of municipal findings. During this study period, and accounting the simultaneous detections in both panels, the average annual number of first reports was 93.5 municipalities for field sampling and 39.4 for citizen science. The Figure 6 describes the evolution of the performance of both surveillance systems, with a Pearson’s correlation coefficient of -0.284 between field and citizen science discoveries (not taking into account simultaneous findings).

Regarding the timing of reciprocal confirmations between strategies, mean data suggest that municipalities initially discovered through citizen science were subsequently confirmed in greater numbers and more quickly than those first identified through field sampling. Of the 335 citizen-science-based discoveries, 151 (45.1%) were later verified in the field—or accepted without field verification—and subsequently appeared in official listings, after an average delay of 2.21 years. Conversely, 331 of the 930 municipalities first detected by field sampling (35.6%) were later confirmed by citizen science, but with a longer average delay of 3.23 years.

### Detection distance for Aedes albopictus by surveillance strategy

This analysis included 1,664 municipalities: 1,262 during the combined period (335 detected through citizen science and 927 through field sampling) plus 402 municipalities detected through field sampling between 2004 and 2013. Analysis of mean distances in the combined period reveals that citizen science yields a significantly broader detection range than field sampling (mean: 21.58 kilometers vs. 11.34 kilometers; Student’s *t*-test, *p* < 0.01). As expected for such a temporally heterogeneous dataset, significant differences were observed across years, as well as an interaction between year and surveilance strategy (ANOVA, *p* < 0.01). Post-hoc Tukey comparisons identified significant differences at the beginning of the period, specifically in 2014 for citizen science and 2015 for both strategies (*p* < 0.05).

When municipalities detected by field sampling prior to 2014 were added, the mean distance did not change significantly, showing only a slight increase to 11.89 kilometers. Exclusion of municipalities with simultaneous detection or due to specific considerations, as exposed, did not affect the statistical significance of the results (data not shown).

The results are depicted in a boxplot in Figure 7 where the horizontal line inside each box represents the median. Due to extreme outliers in the full dataset, a simplified representation is provided retaining values within 1.5 times the interquartile range (IQR).

**Figure 7.**
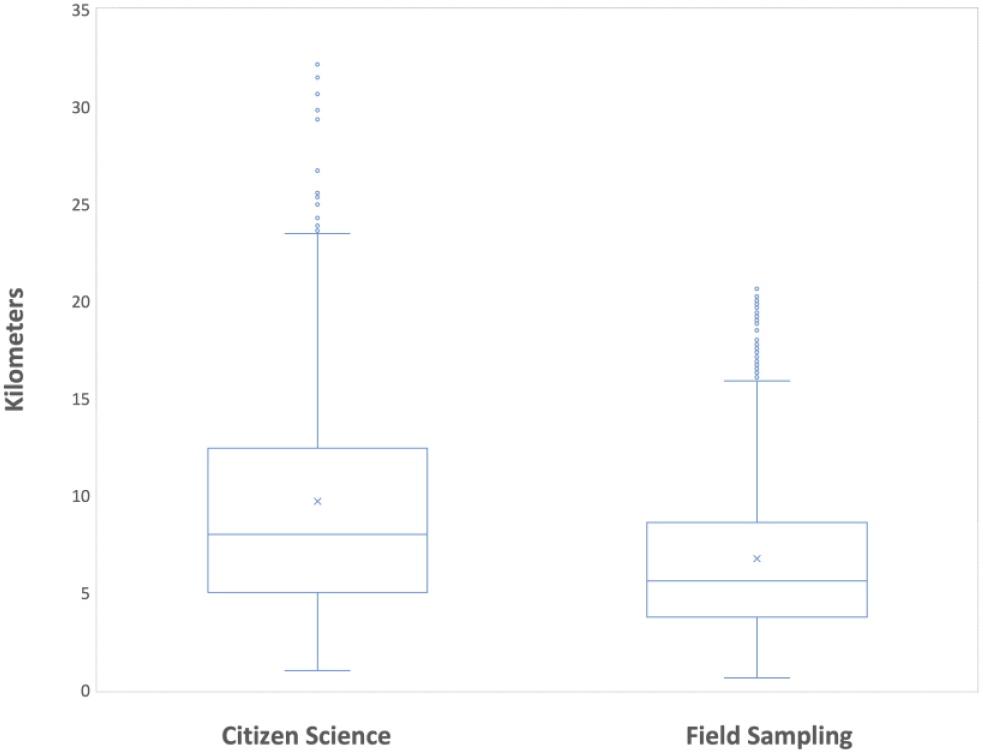
Comparison of distances between detections of Ae. albopictus obtained through citizen science and field sampling (2004–2024) (N=1,476).

Figure 8 displays the distances on the map using the same colour coding as in Figure 5. All positive municipalities are included, with field sampling covering the period 2004–2024, except for previously noted exclusions.

**Fig 8.**
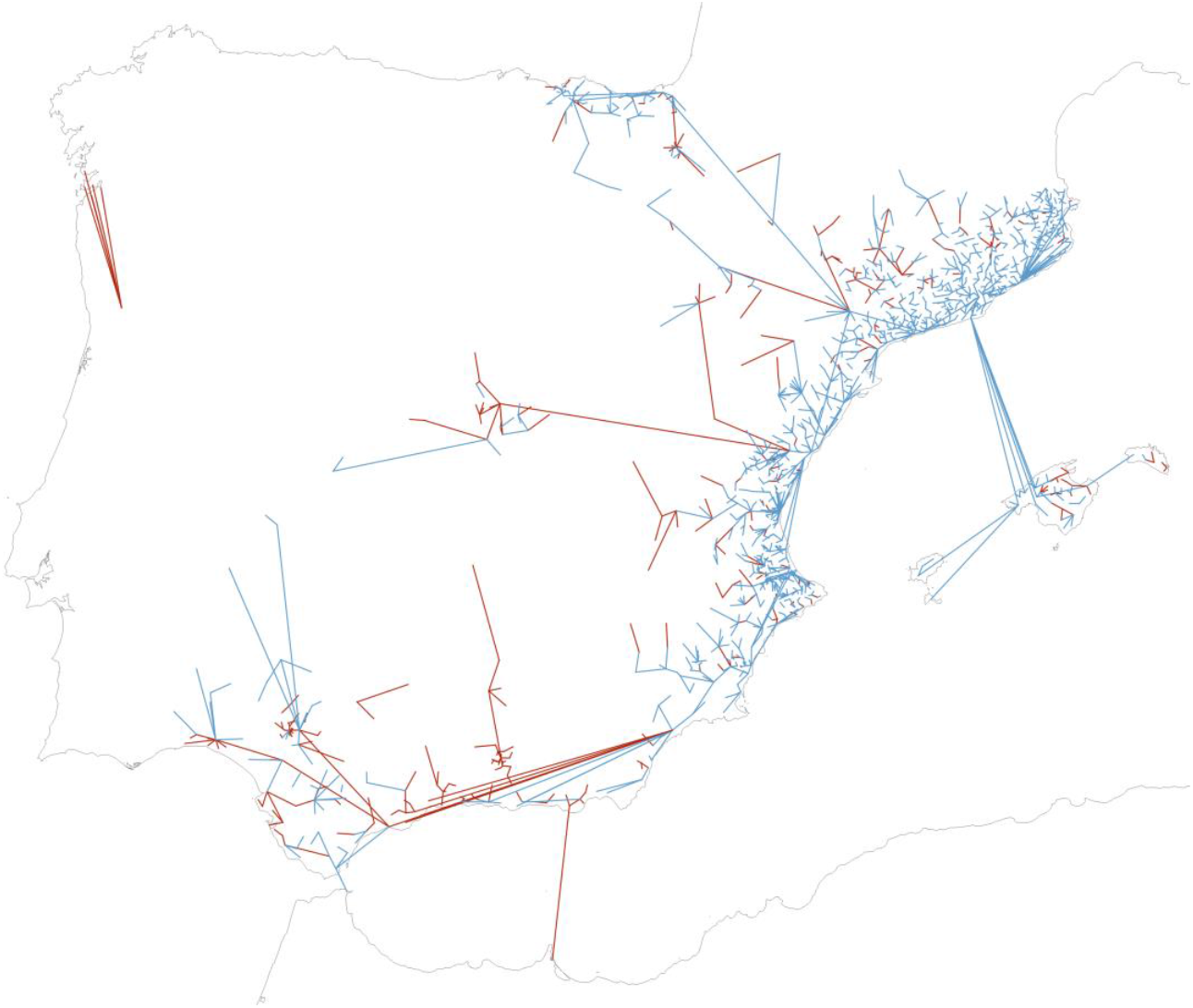
Geographical distances between municipalities at their first detection of Ae. albopictus and the nearest positive municipality for the species up to the previous year, over the entire period 2004–2024 (blue line: new municipality detected by field sampling; red line: new municipality detected by citizen science; simultaneous discoveries not displayed). The Canary Islands are excluded from the analysis and not displayed. Map reference: BDLJE CC-BY 4.0, National Geographic Institute.

### Aedes albopictus detection by citizen science

Of the total 110,939 reports of all categories received by Mosquito Alert from Spain, 56,886 corresponded to adult mosquitoes and were submitted from 2,414 municipalities, covering 29.6% of the total. This corresponds to an average of 23.56 reports per municipality, although with substantial variation due to population distribution and thus, number of users (maximum per municipality: 4,916; minimum: 1; standard deviation: 123.43).

The subset of reports positively identified as *Ae. albopictus* includes 18,398 submissions from 1,091 municipalities, yielding an average of 1.69 reports per municipality per year. However, 287 municipalities received only one report, and in 120 of those cases, that report was the first detection of the species, triggering an alert. Of these 120 municipalities, 43 were later included as positive into the official surveillance listings, implying that from the 335 citizen science discoveries, 77 are currently supported by a single citizen report and -to our knowledge- have not been confirmed through field sampling yet.

Citizen-submitted data has provided the first indications of the presence of *Ae. albopictus* in Castile and Leon (in Sotillo de la Adrada in 2022), Melilla (Melilla in 2023), and Cantabria (Castro-Urdiales in 2024), none of which have been confirmed in the field to our knowledge, which is especially relevant for Melilla as only one citizen report was received. Presence in Castile-La Mancha was also first detected by Mosquito Alert (Illescas in 2015), and currently includes 15 positive municipalities, of which 14 were detected via citizen science, with no field validation neither to our knowledge.

Alongside previously published first detections in Andalusia (Alhaurín de la Torre in 2014) [57], Aragon (Huesca in 2015) [58] and Galicia (Moaña in 2023) [59], citizen contributions through Mosquito Alert have provided the first reports of the species in 7 out of Spain’s 19 ACs.

Early citizen reports were received in the Community of Madrid capital city (2014), as well as from Valdemoro (2015) and Pinto (2016), although they were not confirmed by field sampling, and were deemed as low reliability. In contrast, the region’s field surveillance program officially recorded the first finding in 2016 in Perales de Tajuña [42].

## 4. Discussion

Entomological surveillance in Spain is a regional responsibility, but not a legal obligation. Since 2004, surveillance programs have typically only been implemented once colonization had already reached considerable levels—a situation poorly aligned with the preventive approach required for a rapidly spreading species, particularly in relation to neighbouring regions. To our knowledge, some Autonomous Communities (AC) still do not carry out active field surveillance, meaning that citizen reporting remains the sole detection means across large areas of the country. Nevertheless, cooperation between public services, academia and citizen platforms is advancing rapidly. Currently, the Ministry of Health and six AC with active surveillance programs maintain joint protocols with Mosquito Alert, enabling integrated and rapid responses to new detections, and fostering collaboration in data exchange, public engagement, and vector control. In some of these regions -such as Catalonia and Madrid- citizen detections are routinely used to target field operations.

In contexts lacking historical surveillance, the first detection of an invasive species often raises doubts about the plausibility of its prior absence; contrary to intuitive expectations, a new finding does not necessarily indicate a recent arrival. Such uncertainty should be assessed in light of the historical field sampling efforts when available -as exemplified by the ECDC [37], and, in the case of citizen science data, by estimations of the prior user sampling effort. Otherwise, there is a risk of misinterpreting some findings as species’ expansion when, in reality, we may merely be correcting knowledge gaps about ancient distributions. Occasionally, direct communication with local residents has helped to determine that the species had already been present for some time. This was the case with the first *Ae. albopictus* detection in 2004 (likely present since 2002), with *Ae. japonicus* in 2018 (suspected presence since 2015), and with *Ae. albopictus* in Galicia in 2023 (suspected presence since 2022), based on personal communications.

On the other hand, detection does not necessarily imply establishment, defined as the persistence of reproductive populations for more than a year. While citizen science can provide further evidence of establishment through subsequent repeated reporting, field confirmation requires further sampling, which is often not feasible due to the need to prioritize monitoring uncolonized areas.

This work compiles data on the distribution of the three invasive mosquito species in Spain over the past 21 years, examining both the dynamics of detection processes and spread at the municipal level. While the use of municipalities as base units facilitates practical management, it also adds heterogeneity related to notable differences in surface, population density and also to contagious distribution of urban vectors, impacting detection dynamics as seen in large cities such as Madrid, where a low number of detections in some districts implied listing the whole municipality. Additionally, we did not consider seasonal population fluctuations due to tourist affluence, particularly significant in coastal areas during peak vector activity, considerably increasing the number of people exposed to biting risk.

### Species distribution

As expected, all three species are progressively increasing their presence in Spain since first reporting, likely dispersing through heterogeneous mechanisms and at varying rates. Mosquito Alert detected *Ae. japonicus* in Spain for the first time in 2018 and in subsequent years, citizen science—combined with field sampling—played a crucial role in approximating a more accurate distribution in 111 municipalities. The distribution of *Ae. japonicus* is basically northern in Spain, likely due to its narrower ecological tolerance and preference for deciduous forest habitats [18] in rural, less densely populated environments than *Ae. albopictus* [60]. This species is often detected —like the initial discovery—in remote areas far away from major transport routes or urban areas. Its spread operates through poorly understood mechanisms though it is assumed to rely more on active flight than passive transport, thus highlighting an impressive dispersal capacity. Therefore, and as seen in other countries [61], future findings are likely in regions already silently colonized. Citizen monitoring of *Ae. japonicus* is challenging due to its lower aggressiveness toward humans than of *Ae. albopictus* or *Ae. aegypti*, which typically have rapid public impact upon arrival. It has been suggested that one to three years are necessary for *Ae. japonicus* to gain sufficient visibility after establishment [62].

In the case of *Ae. aegypti*, introductions via maritime transport to the Canary Islands—in addition to those of *Ae. albopictus*—combined to its presence in Cabo Verde and nearby Madeira suggest a high risk of establishment in the archipelago and potential spread to the mainland, given the intensity of trade and travel connections. Integrated control actions by local authorities have achieved eradication successes [55]. Promoting citizen collaboration would be essential in this context, as citizen science can cover entire islands, thereby increasing detection sensitivity beyond the first-tier field surveillance around colonized areas and entry points.

Among the three invasive mosquitoes, the majority of detected positives (1,768 from 1,813) corresponded to *Ae. albopictus*, a species now recorded in 18 of the 19 ACs—Asturias being the sole exception together to Canary Islands when considering eradications. The rapid spread of the Asian tiger mosquito throughout the Mediterranean region from 2004 has been driven by the presence of highspeed transport networks [40], favourable climatic conditions [63] and suitable urban environments: although it is present in less than 22% of the national land, the species impacts more than 66% of the population.

The detection progress of this species followed two different patterns. An initial period with a low but steadily progressing number of new detections per year between 2004 and 2013, affecting mostly Catalonia but also Valencia, Murcia and the Balearic Islands. From 2014 onwards, the inclusion of citizen science coupled to renewed administrative efforts yielded a strong increase in yearly detections, though with a higher heterogeneity. This was probably due to stochasticity, climate, variation in transport activity, increasing number of actors involved in sampling, varying budgets and -for citizen science-fluctuating levels of public promotion. The present colonization rates in these four combined regions average 83% of municipalities, and nearly 100% if considering only coastal areas. In the whole Spain, the number of affected municipalities, surface, and population exposed have all steadily increased over the past 20 years and many areas remain suitable for colonization, moreover considering climate change [63]. A future deceleration in the pace of expansion process could be hypothesized as the species spreads inland using lower transport flows, finding less suitable climates and larger, less populated territories. A geographically stratified permutation test revealed that, in most years, the average population density of newly invaded municipalities was significantly higher than expected under the null model. Likely, this reflects the species’ tendency to first establish in densely populated coastal municipalities before expanding inland into less densely populated regions along time. This pattern suggests a consistent bias toward invasion in denser urban areas within provinces, although possible sampling deviations should also be considered as discussed in the next section.

### Dynamics of surveillance strategies for *Aedes albopictus*

Vector surveillance as presented here relies on two fundamentally different data sources, so any analysis must consider the nature and context of each of them.

Field detection success largely depends on the accuracy of the baseline hypothesis about risk areas driving the planning of the surveys. On the other hand, citizen detections are related to direct mosquito-human interactions -resulting from the degree of perception of insect presence by each participant, balanced by the amount of collective sampling effort. Although this was not used in this work, the Mosquito Alert app provides automatic estimates of the number of users in a given area, allowing to balance the input data with the user dedication. Just like placing more traps for field surveys, this effort may change by increasing promotion in regions where an increased data flow is desired.

In addition to sampling effort random heterogeneity, sampling biases must also be considered in this context, given that both surveillance strategies tend to prioritize large, densely populated municipalities typically containing relevant transport hubs and points of entry, and where most potential citizen contributors are also present. Field sampling might thus be more suited to low population municipalities, where less motivated users are present, and for confirmation of findings; whereas citizen science is perhaps best suited for detections in urban areas and to remote locations.

In this work, citizen contributions accounted for 24.6% of all *Ae. albopictus* detections, preceding field surveillance in 335 municipalities, though the share would increase to 31.8% if including 98 additional municipalities detected in the same year from both data sources. This simultaneous reporting could arise from two scenarios: in some cases, official confirmation may just be the result from the field check of the citizen report -or its acceptance into the official listing without any verification, likely in municipalities adjacent to previously affected ones. Conversely, announcement of a field-based detection can also trigger a rapid surge of positive citizen reports. Unfortunately, in most cases it is not possible to determine the precise sequence of events due to reporting delays, and limited spatio-temporal metadata of field records. Field confirmation should be routinely triggered: to date, 184 out of the 335 citizen-reported positives remain unverified in the field, 77 of them being critical as they were supported by a single report.

Whereas the mean number of discoveries per year are mostly uncorrelated between both data sources, the frequency and time to mutual confirmation on a municipality basis are similar in both directions; although the mean time-to-confirmation is slightly shorter when alerts are originated by citizen science, no conclusions can be drawn due to many undocumented interactions occurring in the detection-verification process, as previously discussed. Convergences and discrepancies between traditional sampling and citizen science further illustrate the complexity of invasive mosquito species detection patterns, and underscore the need for coordinated action protocols between managers of each approach.

### Field sampling

Most detections of new invasive mosquito species in Spain in this study (1,335 cases out of 1,813) have been provided by field sampling, the only surveillance strategy in place for 10 years. Data provided by entomological field operation are irreplaceable due to their authoritative nature as they provide georeferenced, physical evidence coupled to professional taxonomic classification, and are particularly well-suited for proximity-based monitoring.

However, this strategy is constrained by costs and requires specialized personnel, thus challenging wide-area early detection strategies. Results collected by a variety of actors, using non-standardized methods targeting different life stages of the mosquitoes, are often not fully manageable if they lack spatial, temporal, or methodological metadata. As an example, oviposition traps follow variable designs, sizes and substrates, in spite of intents of standardization [39]. Identification methodologies are also not formally standardized, commonly relying on morphological criteria and molecular techniques if available. Additionally, as field surveillance is decentralized, the results are often not published within a timeframe allowing for coordinated responses beyond local scales. In some cases, indirect data collection via questionnaires may add data degradation, corrected here by confrontation to published literature and author’s own data.

### Citizen science

In terms of detections per year, Mosquito Alert achieved an average performance close to 42% of that of field-based surveillance. This is a noteworthy figure, considering the lower costs of this approach and its advantages in scalability, geographic coverage, real time outputs, big data for modelling epidemiological risks and communication capabilities raising awareness and empowering the public to an increasing social challenge in this era of change [64]. Given that Mosquito Alert has not yet benefited from systematic promotion campaigns across Spain, there remains substantial potential for improving performance by expanding the user community.

Since 2023, actionable citizen surveillance in Mosquito Alert has been enhanced by implementing an AI module for image-based species identification, operating autonomously in real time. By combining the geolocation of a suspect report, a species AI scoring threshold and the municipal presence/absence map from this study, the system triggers alerts and send early warning notifications to regional stakeholders. If the score does not meet the threshold the report is forwarded to the experts team, in a slower process that offers, however, higher accuracy and helps with AI’s false negatives.

As discussed, the reach of Mosquito Alert facilitates the detection of events unlikely to be captured by field surveillance, such as long-distance jumps far away from the invasion front. Its broad geographic scope is evident in the analysis of distances between newly detected municipalities and the nearest previous one, which is almost doubling the traditional surveillance (21.58 kilometers versus 11.34 kilometers), consistently with earlier findings for the 2014–2015 period when the respective averages were 37 kilometers and 20 kilometers [65]. This calculation, however, is a highly simplified interpretation of spatial proximity, since closeness by itself does not imply direct spread. Long-distance dispersal—especially in insular settings—may originate from a range of sources, by various dispersal forces mostly driven by human-mediated factors and geographical barriers; multiple historical introductions into the Iberian Peninsula are also suspected [53]. Therefore, our analysis is not intending to describe migration flows, but rather to illustrate the platform’s scalability and sensitivity to remote regions where formal surveillance is often lacking -probably due to that very same perception of remoteness.

Citizen science faces specific challenges, some of them related to the engagement of volunteer communities (both contributors and experts), requiring sustained local promotion commitment to maintain the data flow, which volume is directly correlated to the promotion of the system. A particularly relevant issue is data authentication, stemming from the non-physical nature of citizen contributions needing digital entomology protocols for image classification. Reports with photographs constitute valid evidence when received in sufficient numbers and classified by trained entomologists [66] and a decade of experience has shown such data to reliably indicate the actual presence of the species, reducing the uncertainty by expert consensus triple-blind classification into categories and likelihood levels, combined with AI support. These single reports not confirmed in the field may correspond to recent events, but also to inaccurately geolocated submissions, as the app allows users manual geolocation. The risk of intentional fraud cannot be entirely excluded and is expected to increase as total numbers of participation rise, but is routinely mitigated by manual screening of suspicious images, data consistency protocols, and user confirmation enquiries. These limitations underline the need for integrated confirmation protocols, either through field sampling or subsequent reporting from other users.

Citizen science seeks interoperability: cooperation between Mosquito Alert and other platforms like GLOBE and iNaturalist under the Global Mosquito Alert Consortium (GMAC), allows for integration of citizen data sources in an unified format, easy-to-access citizen data framework (https://www.mosquitodashboard.org/), and GBIF format. These data repositories may help improve upon the current ECDC maps at NUTS 3 level, which offer limited utility for actionable response.

### Emerging surveillance frameworks: managers, experts and citizens

During the second half of the 20^th^ century most concerns over vector-borne diseases in Europe focused mainly on the potential arrival of exotic *Anopheles* single females infected by tropical *Plasmodium* species via air travel, posing a risk of local airport malaria cases. To a large extent, surveillance at airports and other points of entry still today stems from that situation. However, the current vector-related challenges are more complex, involving the displacement of large batches of immature phases of *Aedes* mosquitoes leading to establishment and -coupled to pathogen introductions-making possible eventual local transmission at any time.

The scale-up in surveillance capacity required to respond effectively to a rapidly worsening problem must come through integrated solutions that combine traditional resources with emerging tools. Field entomological surveillance remains essential and irreplaceable, as it has been for decades, but it must now benefit from synergies with new technologies—such as intelligent automatic traps [67,68]—and with strategies rooted in social cooperation and data democracy. Innovation will enable faster and more accurate systems to confront emerging threats, which can no longer be addressed by local sampling and static mappings alone.

The integration of citizen-based surveillance into the interface between society, public administration, the private sector and the environment, expands monitoring capacities at low cost toward new standards. However, such a system works under complex interactions with some extent of operational uncertainty, as external factors such as climate, public funding, and citizen engagement influence the outcomes. This underscores the need to automatically compensate for sampling bias and to establish good indicators of efficiency. A case study of conjoint work between field teams and Mosquito Alert occurred in August 2023, when *Ae. albopictus* was first detected in a report from Galicia. The AI issued a real-time alert that was field-confirmed in less than 72 hours by REGAVIVEC, the official body in charge [59]. Subsequent media outreach and formal promotion of Mosquito Alert did raise by the end of 2024 in Galicia 2,468 citizen-derived reports of mosquitoes, 776 of which described *Ae. albopictus* individuals and resulting in verified detections in six municipalities (plus a seventh not field-confirmed), contributing to delineate the species’ distribution and to surveillance guidance, public communication, and risk analysis in 17 months.

## 5. Conclusions

The distinct characteristics of each data collection strategy, with their respective strengths and limitations, highlight the need for standardization and cooperative integration to enhance overall effectiveness by unifying efforts within a shared, public, accessible, and collaborative system capable of translating results into real-world problem management.

The municipal listing presented here is a consolidated, high-confidence summary of currently available information in spatial and temporal terms, about the invasion process of Spain by *Ae. albopictus, Ae. aegypti*, and *Ae. japonicus* over the 20 years elapsed since the first detection of *Ae. albopictus* in 2004. This compilation has been built upon contributions from both professional entomological sampling, and citizen science data through the Mosquito Alert platform.

This work is a first glance of the capacities of an integrated approach gathering departments of the Ministry of Health, the Ministries for Environment and Livestock, regional governments, municipalities, scientists, stakeholders, private companies and citizenship. Although interactions already exist facilitated by working groups under the umbrella of the Ministry of Health, further steps are needed to formalize and standardize processes in order to develop and stabilize a truly integrated, semiautomated surveillance system. Such a system should guarantee interoperability and accessibility under the FAIR data principles (Findable, Accessible, Interoperable, and Reusable), setting up structured information flows including the public.

To build such a system—overcoming jurisdictional barriers and competency conflicts thus expanding the capacities of previously fragmented schemes—professional One Health networks in entomology -such as VectorNet-ES in coordination with the Ministry of Health would serve as an effective interface. This joint approach, now entering a phase of consolidation, is a pioneering step forward in a state-level collaboration between all actors and an informed, empowered citizenry.

## Author Contributions

Conceptualization, Roger Eritja, Isis Sanpera-Calbet, John Palmer and Frederic Bartumeus; Data curation, Roger Eritja, Isis Sanpera-Calbet, Sarah Delacour-Estrella, Ignacio Ruiz-Arrondo, Mikel Alexander González, Francisco Collantes, María Cruz Calvo-Reyes, Marian Mendoza-García, David Macías-Magro, Pilar Cisneros, Aitor Cevidanes, Eva Frontera, Inés Mato, Fernando Fúster-Lorán, Miguel Domench-Guembe, María Elena Rodríguez-Regadera, Ricard Casanovas-Urgell, Tomás Montalvo, Miguel Ángel Miranda, Jordi Figuerola and Javier Lucientes-Curdi; Formal analysis, Roger Eritja, Isis Sanpera-Calbet, Sarah Delacour-Estrella, Ignacio Ruiz-Arrondo, Maria Àngels Puig, Mikel Bengoa-Paulís, Pedro María Alarcón-Elbal, Carlos Barceló, Simone Mariani, Yasmina Martínez-Barciela, Daniel Bravo-Barriga, Alejandro Polina, José Manuel Pereira-Martínez, Mikel Alexander González, Santi Escartin, Rosario Melero-Alcíbar, Laura Blanco-Sierra, Sergio Magallanes, Francisco Collantes, Martina Ferraguti, María Isabel González-Pérez, Rafael Gutiérrez-López, María Isabel Silva-Torres, Olatz San Sebastián-Mendoza, María Cruz Calvo-Reyes, Marian Mendoza-García, David Macías-Magro, Pilar Cisneros, Aitor Cevidanes, Eva Frontera, Inés Mato, Fernando Fúster-Lorán, Miguel Domench-Guembe, María Elena Rodríguez-Regadera, Ricard Casanovas-Urgell, Tomás Montalvo, Miguel Ángel Miranda, Jordi Figuerola, Javier Lucientes-Curdi, Joan Garriga, John Palmer and Frederic Bartumeus; Investigation, Roger Eritja, Miguel Ángel Miranda, Jordi Figuerola, Javier Lucientes-Curdi and Frederic Bartumeus; Software, Isis Sanpera-Calbet, Joan Garriga and Frederic Bartumeus; Supervision, John Palmer and Frederic Bartumeus; Validation, Roger Eritja, Isis Sanpera-Calbet, Sarah Delacour-Estrella, Ignacio Ruiz-Arrondo, Maria Àngels Puig, Mikel Bengoa-Paulís, Pedro María Alarcón-Elbal, Carlos Barceló, Simone Mariani, Yasmina Martínez-Barciela, Daniel Bravo-Barriga, Alejandro Polina, José Manuel Pereira-Martínez, Mikel Alexander González, Santi Escartin, Rosario Melero-Alcíbar, Laura Blanco-Sierra, Sergio Magallanes, Francisco Collantes, Martina Ferraguti, María Isabel González-Pérez, Rafael Gutiérrez-López, María Isabel Silva-Torres, Olatz San Sebastián-Mendoza, María Cruz Calvo-Reyes, Marian Mendoza-García, David Macías-Magro, Pilar Cisneros, Aitor Cevidanes, Eva Frontera, Inés Mato, Fernando Fúster-Lorán, Miguel Domench-Guembe, María Elena Rodríguez-Regadera, Ricard Casanovas-Urgell and Tomás Montalvo; Writing – original draft, Roger Eritja; Writing – review & editing, Isis Sanpera-Calbet.

## Data Availability Statement

Both the authors and Mosquito Alert hold the copyright of the information. The full dataset is available for download at https://doi.org/10.5281/zenodo.15869763 under a Creative Commons CC-BY NC-SA 4.0 International licensing scheme.

## Acknowledgments

We gratefully acknowledge the contributions of the Mosquito Alert community, who generously volunteered their time and effort to support this project. This includes both the anonymous citizen scientists who submitted reports and the expert community, whose continued efforts in annotating observations and sharing their knowledge have been invaluable.

## Abbreviations

YFV: Yellow fever virus
DENV: dengue virus
CHKV: chikungunya virus
ZIKV: Zika virus
WNV: West Nile Virus
IHR: International Health Regulations
CCAES: Centro de Coordinación de Alertas y emergencias Sanitarias
PNPVC: Plan Nacional de Vigilancia, Prevención y control de las Enfermedades Transmitidas por Vectores
AC: Autonomous Community
ECDC: European Centre for Disease Prevention and Control
EFSA: European Food Safety Authority
AI: Artificial Intelligence

